# More than Words: Neurophysiological Correlates of Semantic Dissimilarity Depend on Comprehension of the Speech Narrative

**DOI:** 10.1101/2020.12.14.422789

**Authors:** Michael P. Broderick, Nathaniel J. Zuk, Andrew J. Anderson, Edmund C. Lalor

**Author notes:** Correspondence: Michael Broderick; Edmund Lalor.

## Abstract

Speech comprehension relies on the ability to understand the meaning of words within a coherent context. Recent studies have attempted to obtain electrophysiological indices of this process by modelling how brain activity is affected by a word’s semantic dissimilarity to preceding words. While the resulting indices appear robust and are strongly modulated by attention, it remains possible that, rather than capturing the contextual understanding of words, they may actually reflect word-to-word changes in semantic content without the need for a narrative-level understanding on the part of the listener. To test this possibility, we recorded EEG from subjects who listened to speech presented in either its original, narrative form, or after scrambling the word order by varying amounts. This manipulation affected the ability of subjects to comprehend the narrative content of the speech, but not the ability to recognize the individual words. Neural indices of semantic understanding and low-level acoustic processing were derived for each scrambling condition using the temporal response function (TRF) approach. Signatures of semantic processing were observed for conditions where speech was unscrambled or minimally scrambled and subjects were able to understand the speech. The same markers were absent for higher levels of scrambling when speech comprehension dropped below chance. In contrast, word recognition remained high and neural measures related to envelope tracking did not vary significantly across the different scrambling conditions. This supports the previous claim that electrophysiological indices based on the semantic dissimilarity of words to their context reflect a listener’s understanding of those words relative to that context. It also highlights the relative insensitivity of neural measures of low-level speech processing to speech comprehension.

## 1. Introduction

Understanding a word’s meaning following the coherent linguistic context in which it appears forms the basis for natural speech comprehension. Recent studies have attempted to obtain electrophysiological indices of this process by modelling how brain responses are affected by a word’s meaning relative to its context (Broderick, Anderson, Di Liberto, Crosse, & Lalor, 2018; Dijkstra, Desain, & Farquhar, 2020; Frank & Willems, 2017). One particular approach involved regressing electroencephalographic responses to natural speech against the semantic dissimilarity of words to their preceding context (Broderick et al., 2018). This produced neurophysiological model measures that appeared to be exquisitely sensitive to a listener’s speech understanding. This sensitivity was demonstrated by comparing the neural measures for forward speech and time-reversed speech; by investigating speech processing under different levels of perceived noise; and in tasks where speech was either attended or unattended (Broderick et al., 2018). In each of these experiments, speech understanding coincided with a neural model measure closely resembling the N400 component of the event related potential, which has long been associated with the processing of meaning (Kutas & Federmeier, 2011; Kutas & Hillyard, 1980).

We have proposed that this neural model measure – known as a semantic temporal response function (TRF) – reflects the semantic processing of words in their context. However, given that the approach is based on calculating the semantic dissimilarity of words to their preceding context, it remains possible that the measure reflects a sensitivity to word-to-word changes in semantic content without the need for a narrative-level understanding on the part of the listener. Furthermore, the human brain processes speech at multiple linguistic levels, including the analysis of words’ phonological, lexical, and syntactic properties (Davis & Johnsrude, 2003; de Heer, Huth, Griffiths, Gallant, & Theunissen, 2017; Hickok & Poeppel, 2007; Poeppel, Emmorey, Hickok, & Pylkkänen, 2012; Price, 2010). As such, it is possible that the semantic TRF could also reflect processing at some level lower than semantics, where words are recognised but their true underlying meaning isn’t processed in relation to previous context. Indeed, as mentioned, the morphology and topographical distribution of the semantic TRF shared common characteristics with the N400. And the N400 component has been shown not only to be sensitive to semantic properties of words in context but also to lower-level lexical properties like word frequency and neighbourhood density (Kutas, 1993). Therefore, we wished to further examine the semantic TRF’s sensitivity to the processing of intelligible speech where all words were lexically identifiable, but the overall meaning of the speech narrative was not understood. In other words, we wanted to determine whether the recently derived semantic TRF was specifically sensitive to the processing of words following a coherent, predictive context, thus reflecting semantic levels of processing, or whether it could be elicited by the same speech where word order, and thus context, had been manipulated.

To manipulate the narrative coherence of our stimuli while still presenting each word with perfect intelligibility, we systematically randomised the order of words for a piece of narrative text to create 5 different levels of randomisation, or scrambling. This text was then converted to a speech signal and presented to subjects while their EEG was recorded. Similar forms of linguistic degradation have been used in previous studies to investigate syntactic and combinatorial semantic processing (Humphries, Binder, Medler, & Liebenthal, 2006; Mollica et al., 2018). Inspired by previous studies, where local temporal information in the acoustic (Kiss, Cristescu, Fink, & Wittmann, 2008; Saberi & Perrott, 1999) or linguistic (Lerner, Honey, Silbert, & Hasson, 2011) speech stream was manipulated at gradually increasing temporal windows, we varied the number of consecutive words that could be scrambled to gradually increase the comprehension difficulty of the speech signal. This created graded levels of speech understanding with which we could test the sensitivity of the recently derived measures of semantic processing.

We additionally wished to test the impact of speech comprehension on neural responses relating to the speech envelope. The amplitude envelope of the speech waveform has been highly emphasised in the literature (Luo & Poeppel, 2007; Obleser, Herrmann, & Henry, 2012) and is an important cue for speech perception (Drullman, Festen, & Plomp, 1994a, 1994b; Shannon, Zeng, Kamath, Wygonski, & Ekelid, 1995). This has led researchers to use neural indices of envelope tracking as dependent measures of speech comprehension (Etard & Reichenbach, 2019; Verschueren, Somers, & Francart, 2019). However, reliable envelope tracking has also been shown for an array of signals that do not allow comprehension (Doelling & Poeppel, 2015; Howard & Poeppel, 2010; Lalor, Power, Reilly, & Foxe, 2009; Peña & Melloni, 2012). Fewer studies have compared envelope tracking of speech that is entirely recognisable to the listener but varies in its degree of semantic comprehensibility. Here we do so by deriving TRFs to the speech envelope across our different scrambling levels, allowing us to compare semantic and envelope TRFs for the same speech and the same subjects.

## 2. Methods

### 2.1 Subjects

15 subjects (7 female) aged between 19 and 29 participated in the study. All participants were native English speakers, had self-reported normal hearing, were free of neurological diseases, and provided written informed consent. All procedures were undertaken in accordance with the Declaration of Helsinki and were approved by the Ethics Committees of the School of Psychology at Trinity College Dublin, and the Health Sciences Faculty at Trinity College Dublin.

### 2.2 Stimuli and Experimental Procedure

Stimuli were acquired from a children’s novel (White, 1951). Text from chapters 3-10 of the novel were split into 60 segments corresponding to the 60 trials in the experiment. Each segment then underwent a scrambling procedure to generate 5 different versions. This was done by grouping consecutive words into windows of length *w* and randomly permuting the words within each window. The window lengths were selected as *w* = 1 (unscrambled), 2, 4, 7, 11 so that the interval between window lengths increased by 1 with each *w* (i.e. 2 – 1 = 1, 4 – 2 =2, 7 – 4 =3 etc.). Audio versions of each text segment (including the unscrambled text segments) were then generated using Google’s Text-to-Speech. This software is powered by WaveNet (Oord et al., 2016), a generative model for raw audio based on a deep neural network. It generates realistic human-like sounding voices and has been rated significantly more natural sounding than the best parametric and concatenative systems for English (Oord et al., 2016). Each speech segment, or trial, lasted ~60 seconds and could be heard as one of the five scrambled versions.

Two sets of questionnaires were generated for each trial. The first was a comprehension questionnaire that consisted of 6 questions with 2 possible answers, pertaining to the content of the trial. The second was a lexical identification questionnaire. 6 words were presented with yes/no choices as to whether the word appeared in the trial. 2-4 of these words were nouns/verbs which had appeared in the trial, and the remainder were nouns/verbs which were randomly selected from the rest of the text and did not appear in the trial. No subject reported listening to or reading the novel within at least 3 years of participating in the study.

For the EEG experiment, trials were presented chronological to the story in 12 blocks of 5 trials. Each block contained one trial from each of the 5 different scrambling conditions. Condition order was randomised for each subject and for each block, so that, for example, subject 1 might hear trial 1 with a scrambling window of 11 whereas subject 2 might hear trial 1 with a scrambling window of 4. After each trial subjects were presented with a comprehension and lexical identification questionnaire. Subjects were encouraged to take breaks when needed between trials. Stimuli were presented diotically at a sampling rate of 44.1 kHz using HD650 headphones (Sennheiser) and Presentation software (Neurobehavioural Systems). Testing was performed in a dark, sound-attenuated room, and subjects were instructed to maintain visual fixation on a crosshair centred on the screen for the duration of each trial, and to minimize eye blinking and all other motor activities.

### 2.3 Data Acquisition and Preprocessing

128-channel EEG data were acquired at a rate of 512 Hz using an ActiveTwo system (BioSemi). Offline, the data were downsampled to 128Hz and bandpass filtered between 0.5 and 8Hz using a zero-phase shift Butterworth 4^th^ order filter. To identify channels with excessive noise, the standard deviation of the time series of each channel was compared with that of the surrounding channels. For each trial, a channel was identified as noisy if its standard deviation was more than 2.5 times the mean standard deviation of all other channels or less than the mean standard deviation of all other channels divided by 2.5. Channels contaminated by noise were recalculated by spline interpolating the surrounding clean channels. Independent component analysis (Hyvarinen, 1999) was then performed on the EEG data in order to remove eye blinks. EEG data was transformed into component space and components relating to eye-blinks were identified based on their topographical distribution and component time-series. These components were removed, and the data were then transformed back to EEG channel space. Data were then referenced to the average of the 2 mastoid channels and were normalized to zero mean and unit SD (z units).

### 2.4 Stimulus Characterisation

We wished to test the effect of scrambling condition on the neural encoding of low-level and high-level properties of the speech signal. For the low-level representation we chose the speech envelope, an approach that has been widely used in recent years (e.g., (Kubanek, Brunner, Gunduz, Poeppel, & Schalk, 2013; Lalor & Foxe, 2010). We calculated it by taking the absolute value of the Hilbert transform of the broadband (80 to 3000Hz) speech signal, and passing that through a zero phase-shift low pass filter with a cutoff at 30 Hz.

We represented the context-based meaning of the words in the speech using semantic dissimilarity. Semantic dissimilarity quantifies the semantic relationship between words and their previous context. GloVe – a word embedding model (GloVe; (Pennington, Socher, & Manning, 2014)) - was used to represent each content word in the stimulus as a vector or coordinate in high dimensional space. Vectors were derived by factorizing the word co-occurrence matrix of a large text corpus - in this case Common Crawl (https://commoncrawl.org/). The output is a 300-dimensional vector for each word, where each dimension can be thought to reflect some latent linguistic context. A word’s semantic dissimilarity was estimated as 1 minus the Pearson’s correlation of the word’s vector and the average vector of words from the preceding context. Previous studies have chosen this preceding context to be all previous words in the same sentence (Broderick et al., 2018; Broderick, Di Liberto, Anderson, Rofes, & Lalor, 2020). However, given that, for scrambled speech, sentence boundaries were randomised, we chose to instead estimate semantic dissimilarity by comparing a word vector with the averaged vector of its 10 preceding words. This window was chosen somewhat arbitrarily, however, choosing different context window lengths of 5, 8 and 12 did not qualitatively alter the overall results. The semantic dissimilarity measure was quantified as a vector of impulses, the same length as the presented trial, with impulses at the onset of each content word whose heights was scaled according to their semantic dissimilarity value. Finally, we included an additional feature of word onset as input to the TRF. This is an impulse vector, with impulses at the beginning of each content word, whose height is a constant value of the average of the semantic dissimilarity values in the same trial. The purpose of including this feature was to try to soak up additional variance in the EEG response related to the acoustic processing of the word onset.

Speech waveforms for each trial and each scrambling condition (60 trials x 5 conditions) were generated first (using WaveNet) before the speech envelope or semantic dissimilarity features were estimated.

### 2.5 TRF Estimation and Evaluation

To test the encoding of the lower and higher-level properties of the speech signal we used the temporal response function (TRF). The TRF estimates a linear mapping between some continuous speech feature(s), S(t), and the continuous neural response, R(t).

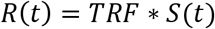

Where ‘*’ represents the convolution operator. S(t) can be a single speech feature, i.e. univariate, or multiple speech features, i.e. multivariate. The TRF for each feature is calculated over a series of time lags between the stimulus and the response, producing a set of temporal TRF weights for each EEG channel. To estimate the TRF we used ridge regression (see Crosse et al., 2016 for details on the TRF)

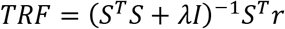

where λ is a regularization parameter that controls for overfitting. A range of TRFs were constructed using different λ values between 0.1 and 1000. The λ value corresponding to the TRF that produced the highest EEG prediction accuracy (see below), averaged across trials and channels, was selected as the regularisation parameter for all trials per subject. TRFs were estimated separately for each scrambling condition by grouping trials from each condition (12 trials per condition) together. Separate envelope TRFs and semantic dissimilarity TRFs were estimated using the envelope and semantic dissimilarity speech representations, respectively. For the semantic dissimilarity TRF, a range of time-lags from 0-700ms was selected (although see below). For the envelope TRF, the time-lag range was selected between -100 and 400ms.

TRFs were evaluated in two ways. First, the TRF weights themselves, across time and across channels, were used as dependent measures. And, second, we used a leave-one-out, cross validation analysis to predict unseen EEG using TRFs fit on other data. The type of approach – known as a forward encoding model – is commonly used to assess whether or not brain responses are sensitive to specific stimulus features (Naselaris, Kay, Nishimoto, & Gallant, 2011). Specifically, for each condition and speech representation, we fit TRFs on 11 out of 12 trials and used those TRFs to predict the EEG data of the held-out trial. And we rotated which trial was held out until all the data had been used. Predicted EEG was compared with the actual recorded EEG data to obtain prediction accuracy, measured using Pearson’s correlation. For the semantic dissimilarity TRF some additional steps were implemented in order to directly test how well semantic dissimilarity captured the semantic processing of words relative to their context. First, TRFs were trained and tested on a narrower window of time-lags, from 200-600ms, rather than the full 0-700ms time-lag window. This was done to ensure that EEG prediction accuracy reflected only neural activity relating to higher level linguistic processing. Although the plotted TRFs (figure 2) show the entire time-lag window (0-700), prediction accuracies (figure 3) were obtained from this narrower window. Second, we wished to establish a stringent baseline against which EEG prediction accuracies from the semantic dissimilarity TRF could be tested. For each stimulus trial and scrambling condition, 5 null semantic dissimilarity representations were generated. This was done by randomly permuting the heights of the impulses in the semantic dissimilarity vectors. In the testing phase of the cross-validation procedure, trained TRFs were used to predict EEG based on the 5 null representations for every recorded trial. The predicted EEG of these 5 null representations was then compared to the actual EEG with Pearson’s correlation. The EEG prediction accuracies of these null representations formed a baseline against which the accuracy of the true representation was compared. Specifically, true EEG prediction accuracy was then calculated by subtracting the mean prediction accuracies from the 5 null representations from the prediction accuracy obtained for the true semantic dissimilarity representation. EEG prediction accuracy for semantic dissimilarity, therefore, refers to the prediction accuracy *difference* between the true speech feature and the average of the 5 null speech feature representations.

### 2.6 Statistical Analysis

For the comprehension and lexical identification questionnaires, above-chance performance was estimated using the binomial inverse cumulative distribution function in MATLAB (Combrisson & Jerbi, 2015), which determines the 95% confidence interval for exceeding chance level based on the number of trials (n = 60) and significance threshold (α = 0.05). Comprehension and lexical identification scores and EEG prediction accuracies were compared using one-way ANOVAs. TRF waveforms (i.e., weights) were tested as being significantly less than zero using a running t-test across subjects. The resulting p values were FDR corrected (Benjamini & Hochberg, 1995). Average TRF waveform differences were measured using a running ANOVA, where TRF weights at each time lag were compared across scrambling conditions. This produced F-values as a function of time-lag. P values corresponding to the running F-values were FDR corrected.

## 3. Results

### 3.1 Scrambling window impacts comprehension despite stability of attention

Subjects were required to answer comprehension and lexical identification questionnaires after each experimental trial. The purpose of these questionnaires was to test subject’s understanding of the speech segment and attention levels during each trial. Figure 1 shows the comprehension and lexical identification questionnaire scores for each condition. Comprehension accuracy was significantly greater than the 95% confidence interval exceeding chance across subjects for scrambling windows of 1 (unscrambled; t_14_ = 13.6, p = 8.94 x 10^-9^) and 2 (t_14_ = 4.2, p = 2.3 x 10^-3^), but not for windows of 4 (t_14_ = 0.78, p = 0.54), 7 (t_14_ = -0.3, p = 0.77) or 11 (t_14_ = -0.86, p = 0.55; one-sample t-test, corrected for multiple comparisons). While the above comparison was designed to test if the group was doing better than a chance level determined by the number trials each subject undertook, we also directly compared the group against 50% (i.e., chance) for completeness. Population performance was significantly greater than chance level (50%) for all scrambling windows (1: t_14_ = 18.8, p = 1.2 x 10^-10^, 2: t_14_ = 7.8, p = 4.6 x 10^-6^, 4: t_14_ = 3.9, p = 2.1 x 10^-3^, 7: t_14_ = 3.5, p = 3.3 x 10^-3^, 11: t_14_ = 3.9, p = 2.1 x 10^-3^; one-sample t-test, corrected for multiple comparisons). A one-way ANOVA revealed a main effect of scrambling condition on comprehension question score; F(4, 70) = 20.79, p = 2.5 x 10^-11^, one-way ANOVA. Pairwise comparisons using Tukey’s Honestly Significant Difference (HSD) procedure indicated that scores for a scrambling window of 1 were significantly higher compared to all other scrambling windows (p < 0.05). Scores for a scrambling window of 2 were significantly higher for windows of 7 and 11 (p < 0.05) but not for 4 (p = 0.1). Comprehension scores for all other windows were not significantly different from each other (p > 0.05).

**Figure 1.**
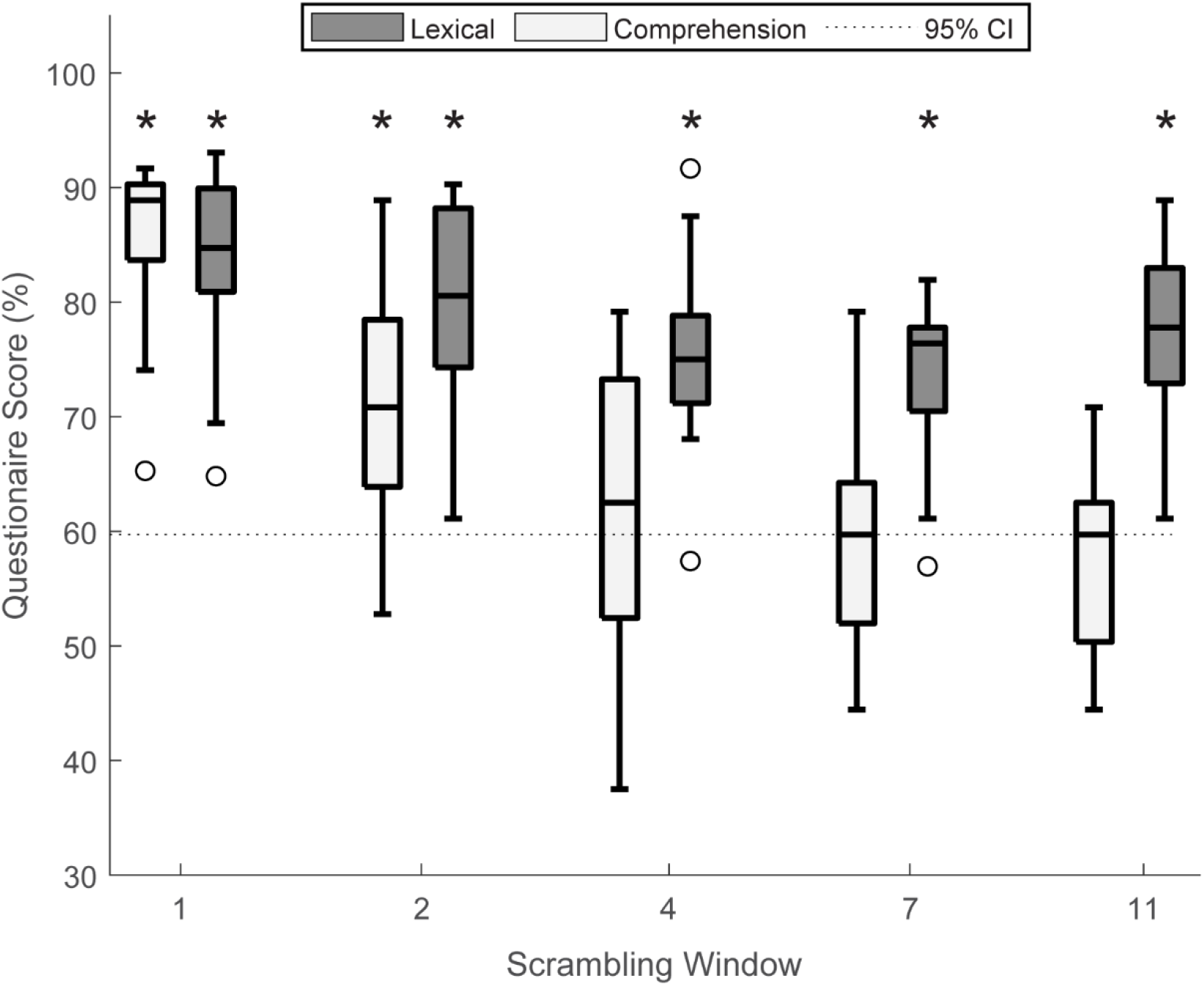
Behavioural effects for each scrambling condition. Box plots indicate comprehension (light) and lexical identification (dark) questionnaire scores for each scrambling window. The 95% confidence interval (CI) exceeding chance level accuracy is indicated by the dashed horizontal line. Stars indicate the scrambling condition where average subject score was significantly above the 95% CI. Lexical identification scores were significantly greater than the 95% CI across subjects for all scrambling windows. Conversely only scrambling windows of 1 and 2 had comprehension scores that were significantly above chance across subjects. Boxplots show median, and first and third quartiles. Whiskers show 1.5 × IQR. Dots indicate outliers.

In contrast to the fall off in comprehension with increased scrambling window length, lexical identification accuracy was significantly greater than the 95% confidence interval exceeding chance for all windows (1: t_14_ = 11.6, p = 7.4 x 10^-8^, 2: t_14_ = 9.1, p = 6.6 x 10^-7^, 4: t_14_ = 7.4, p = 3.2 x 10^-6^, 7: t_14_ = 7.6, p = 3.2 x 10^-6^, 11: t_14_ = 8.9, p = 6.6 x 10^-7^; one-sample t-test, corrected for multiple comparisons) and for chance level performance (1: t_14_ = 16.2, p = 9.2 x 10^-10^, 2: t_14_ = 13.5, p = 3.4 x 10^-9^, 4: t_14_ = 12.7, p = 5.8 x 10^-9^, 7: t_14_ = 12.7, p = 5.8 x 10^-9^, 11: t_14_ = 13.9, p = 3.4 x 10^-9^; one-sample t-test, corrected for multiple comparisons). These behavioural results suggest that, although subjects failed to understand at higher scrambling windows, they remained engaged with the stimulus and were able to successfully identify words that appeared in the trial. There was an additional effect of scrambling condition on lexical identification questionnaire score; F(4, 70) = 3.6, p = 0.01, one-way ANOVA. Pairwise comparisons revealed significant differences between a scrambling window of 1 and scrambling windows of 4 and 7 (p < 0.05). There were no other significant differences between scrambling conditions (p > 0.05).

### 3.2 Semantic TRF weights pattern with comprehension

We next tested whether the semantic dissimilarity TRF was sensitive to a subject’s understanding of the speech narrative they were hearing or whether the measure reflected the more temporally local, low-level processing of word-to-word changes in semantic content. We first examined the weights of the TRF over time-lags, averaged over parietal channels. Figure 2A-B show the time course of TRF weights. TRFs relating to conditions with less scrambling (w = 1 and 2) showed a morphology highly consistent with previous studies investigating the semantic dissimilarity measure (Broderick et al., 2018; Broderick et al., 2020). Weights for these scrambling windows (W = 1 and 2) were significantly greater than zero across subjects for a range of time lags between 400 and 600 ms (Fig. 2B horizontal line, running one-sample t-test, FDR corrected). No such significant difference was found for scrambling windows of W = 4, 7, or 11 (Fig 2B). A running one-way ANOVA was conducted on each time-lag, comparing TRF weights across scrambling conditions. Plotted below figure 2A is the time series of F-values corresponding to each comparison. TRF weights were significantly different across scrambling conditions between 400 and 500ms (p<0.01). However, these F-values did not survive significance testing after correcting for multiple comparisons (0.05<p<0.1). The topographical distribution of the TRF weights averaged over later time-lags (300-600ms; Figure 2C) further reveal responses that shared similar characteristics of the N400 component. Crucially, this pattern of responses was weaker or absent for TRFs derived from conditions with higher word scrambling. This indicates that semantic dissimilarity TRFs do not simply show responses to the semantic difference between a word and previously heard words, but that they rely on the subject’s ability to relate a word to its narrative context.

**Figure 2.**
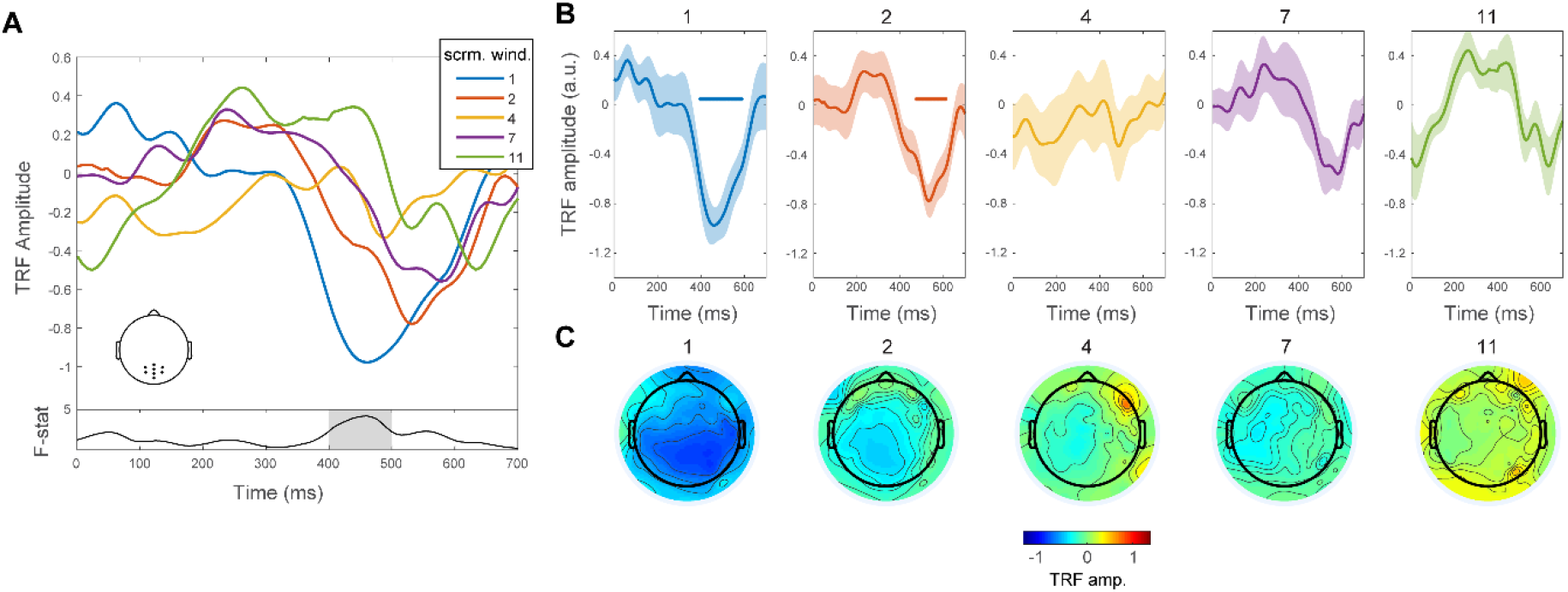
Semantic dissimilarity TRFs for speech at each scrambling window. **(A)** TRF weights averaged over parietal electrodes and across subjects for each scrambling condition. Inset topographical plot shows the selected parietal electrodes Plotted below are the F-values of a running one-way ANOVA, comparing weights across conditions at each individual time-lag. Significant differences across scrambling conditions were observed over time-lags between 400 and 500ms (shaded region; p < 0.01; not corrected for multiple comparisons) **(B)** The same TRF weights plotted in A in separate plotting windows. Horizontal lines in each window indicate the time points where TRF weights were significantly different from zero (FDR-corrected). **(C)** Topographical plots show weight values averaged over later time-lags (300-600ms) and across subjects.

Next, we tested how well TRFs trained on data for each scrambling condition could predict neural responses from held out trials, above a permuted null baseline. Figure 3 shows EEG prediction accuracies averaged over the same set of parietal electrodes for each scrambling window. Below the box plots are topographical plots showing prediction accuracy, averaged across subjects, for each individual channel. The unscrambled speech condition was the only one in which trained TRFs could accurately predict EEG above the null baseline (t_14_ = 3.32, p = 0.025; one-sample t-test, corrected for multiple comparisons). Although the TRF weights for a scrambling window of 2 were significantly greater than zero over later time-lags (Figure 2B), they were unable to significantly predict EEG above the null baseline (t_14_ = 0.67 p = 0.71; one-sample t-test, corrected for multiple comparisons). EEG prediction accuracies for the other scrambling conditions were also not significantly greater than the null baseline (4: t_14_ = 0.43, p = 0.71; 7: t_14_ = 1.7 p = 0.26; 11: t_14_ = -0.37 p = 0.71; one-sample t-test, corrected for multiple comparisons). A one-way ANOVA was used to test the difference in prediction accuracy between scrambling conditions where a significant effect was found; F(4, 70) = 3.83, p = 0.007. Pairwise comparisons using Tukey’s HSD procedure indicated that prediction accuracy for a scrambling window of 1 was significantly higher compared to a scrambling window of 2, 4 and 11 (p < 0.05) but not 7 (p = 0.17). There were no other significant differences between scrambling conditions (p > 0.05).

**Figure 3.**
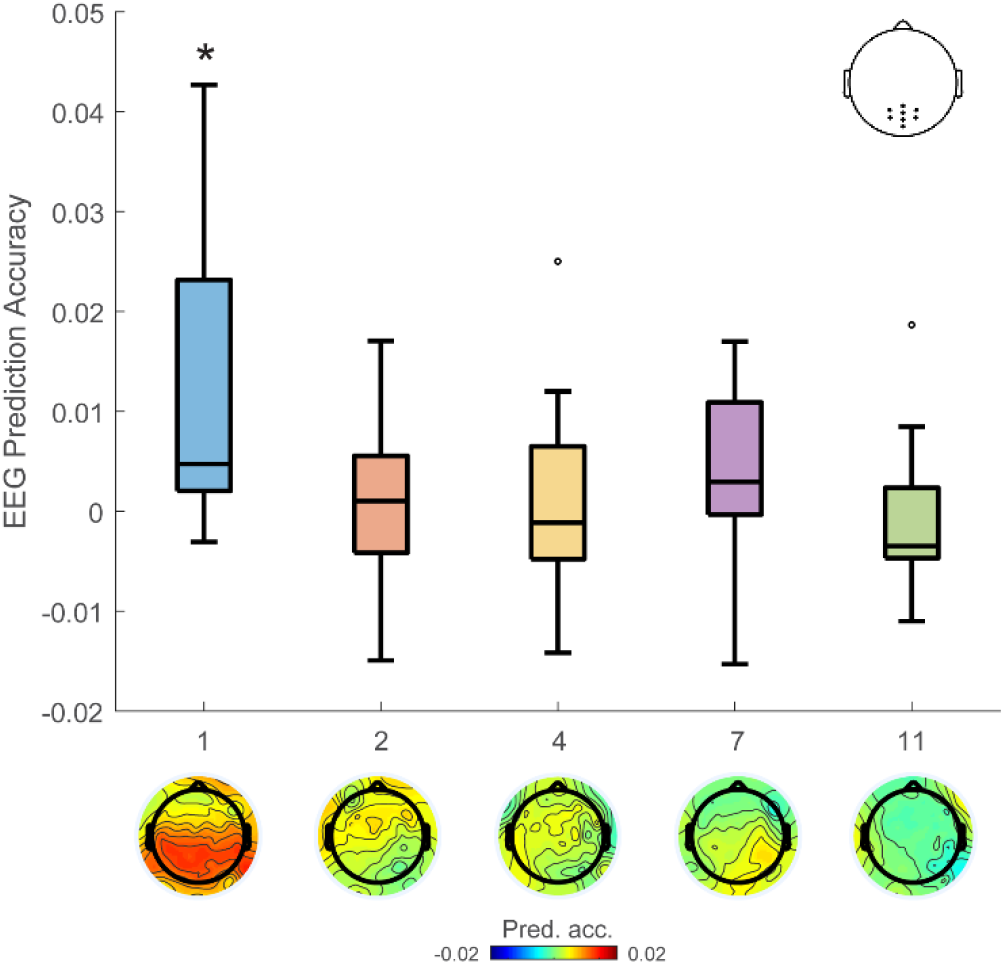
EEG prediction accuracy for semantic dissimilarity. Prediction accuracy was estimated as a trained TRF’s ability to predict EEG from a true semantic dissimilarity representation over its ability to predict EEG for null semantic dissimilarity representation. EEG prediction accuracies averaged over parietal electrodes are given for each scrambling window (1, 2, 4, 7, 11). Asterisk indicates prediction accuracies that were significantly greater than zero. Below are topographical plots of prediction accuracies at each channel averaged across subjects for each scrambling window.

### 3.3 Envelope tracking is unaffected by speech understanding

We tested whether the envelope-based TRF would show a similar sensitivity to scrambling condition and, hence, also reflect narrative speech understanding. Figure 4A shows the TRF weights averaged over temporal electrodes and across subjects for each scrambling window. The weights show a morphology characteristic of speech envelope-based TRFs from previous studies (Di Liberto, O’Sullivan, & Lalor, 2015; Lalor & Foxe, 2010). Unlike the semantic dissimilarity TRF weights, the morphology of TRF weights for all scrambling conditions shared common characteristics. As before, a running one-way ANOVA was conducted on each time time-lag separately, comparing envelope based TRF weights across scrambling conditions. Plotted below figure 4A is the time series of F-values corresponding to each comparison. TRF weights were significantly different across scrambling conditions in a short window between 200 and 220ms (p < 0.05). These F-values did not survive significance testing after correcting for multiple comparisons (p > 0.05).

**Figure 4.**
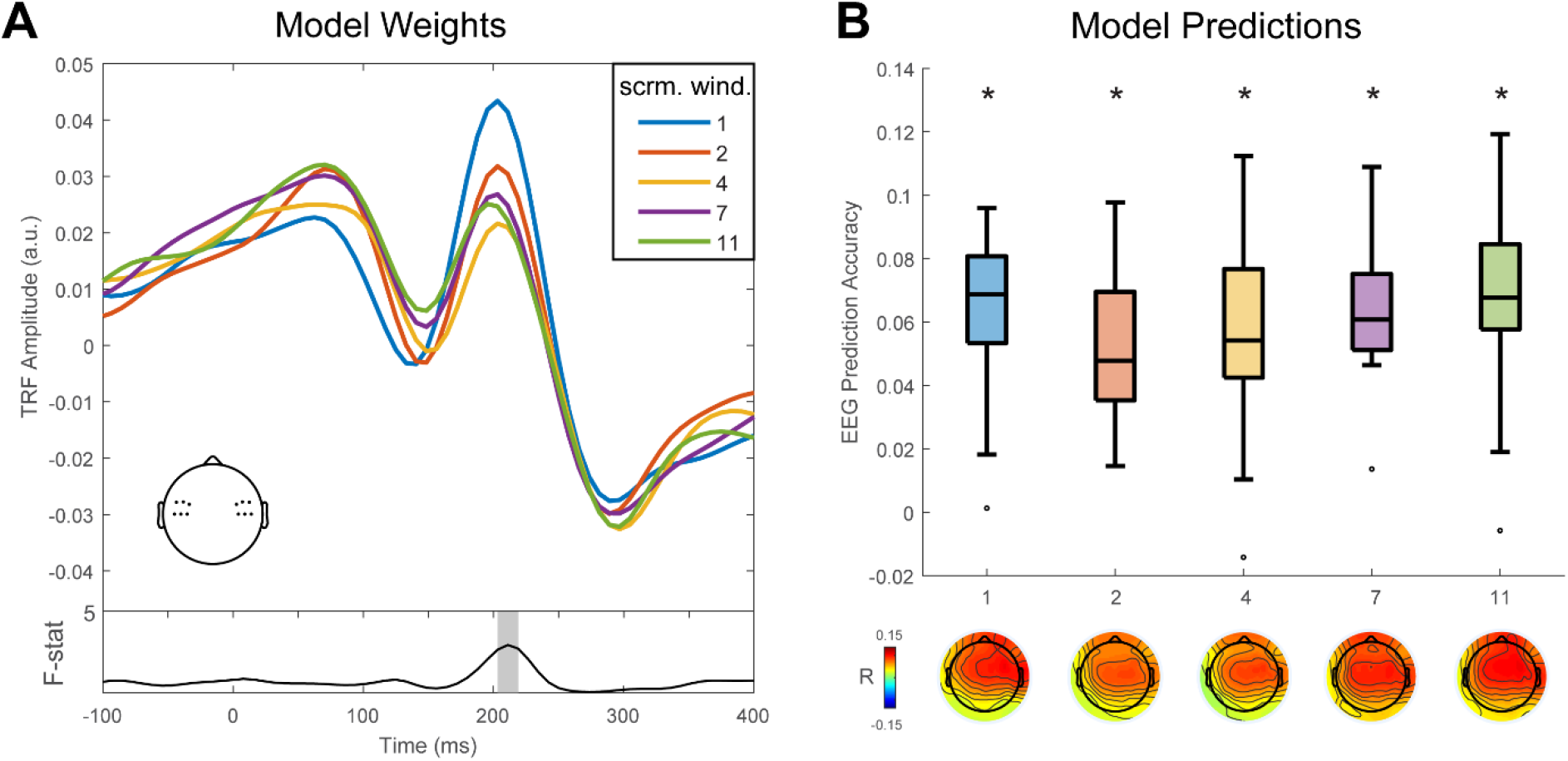
Envelope TRFs for speech at each scrambling window. **(A)** TRF weights averaged over temporal electrodes and across subjects for each scrambling condition. Overlaid topographical plot shows the selected temporal electrodes. Plotted below is the F-values of a running one-way ANOVA, comparing weights across conditions at each individual time-lag. Shaded regions indicate significant (uncorrected F-values). **(B)** EEG prediction accuracies for TRF trained on the envelope. Asterisk indicates prediction accuracies that were significantly greater than zero. Below are topographical plots of prediction accuracies at each channel averaged across subjects for each scrambling window.

We tested each trained TRF’s ability to predict unseen EEG. Figure 4B shows prediction accuracy, averaged across temporal electrodes. Below the boxplots shows the topographical distribution of prediction accuracies, average across subjects. TRFs from all 5 scrambling conditions were able to significantly predict EEG (1: t_14_ = 8.9, p = 8.2 x 10^-7^, 2: t_14_ = 8.7, p = 8.2 x 10^-7^, 4: t_14_ = 6.2, p = 2.2 x 10^-5^, 7: t_14_ = 10.9, p = 1.6 x 10^-7^, 11: t_14_ = 8.4, p = 9.9 x 10^-7^; one-sample t-test, corrected for multiple comparisons). Differences in prediction accuracy between scrambling conditions were tested using a one-way ANOVA. No significant differences were found for mean prediction accuracy across scrambling conditions; F(4, 70) = 0.8, p = 0.53. This indicates that, although subjects could not grasp the meaning of the speech they were hearing, their EEG still reliably tracked the acoustic properties of the speaker.

EEG prediction accuracies of envelope based TRF were estimated based on a broad window of timelags (-100 to 400ms). However, we observed small differences between scrambling conditions at peaks in the TRF for later time-lags (200-220ms). Therefore, it is possible that this window could represent some locus of comprehension and, thus, training and testing TRFs in this window might reveal differences in EEG prediction accuracies relating to speech understanding. We trained envelope TRF using a time-lag window centred around the P2 peak (175 to 225ms) and found that this was not the case. Prediction accuracies for all scrambling conditions were reduced but significantly greater than zero (p < 0.05; corrected for multiple comparisons). There was no significant effect of scrambling condition on EEG prediction accuracy; F(4,70) = 1.7, p =0.16, one-way ANOVA.

### 3.4 Semantic Dissimilarity TRF predicts comprehension scores

Finally, we wished to test whether the TRF measures could reliably predict an individual’s comprehension score for each scrambling condition. As seen above, the averaged weights and prediction accuracies for the semantic dissimilarity TRF indeed reveal responses that patterned with scrambling condition. However, we wished to examine whether comprehension scores at the level of individual subjects and individual trials could be predicted using our neural measures. We therefore generated linear mixed effect (LME) models, which took prediction accuracies from the semantic dissimilarity based TRFs and envelope based TRFs as independent variables and comprehension scores as dependent variables, with random effects of individual subjects. One of the LME models predicted comprehension scores averaged across trials for each scrambling condition and the other predicted comprehension scores at the level of individual trials. Predictor variables were normalized before being inputted to the model. Table 1 shows the predictor variable coefficients for the first model. There was a significant main effect of average semantic dissimilarity prediction accuracy on comprehension score (t = 2.93, p = 0.005) and no main effect of average envelope prediction accuracy (t = - 0.76, p = 0.44). Furthermore, we tested the interaction between predictor variables which had no effect on comprehension score (t = - 0.39, p = 0.7)

**Table 1.** Subjects LME Model. Model weights for the linear mixed effects model predicting comprehension scores at the level of subjects

|  | Coefficient | S.E. | t statistic | p |
| --- | --- | --- | --- | --- |
| **Sem. Dissim.** | **4.62** | **1.59** | **2.93** | **0.005** |
| Env. | - 2.1 | 2.75 | - 0.76 | 0.44 |
| Sem. Dissim. * Env. | - 0.56 | 1.45 | - 0.39 | 0.7 |

Next, we looked at the TRF output measures of individual trials and their ability to predict comprehension success. Table 2 shows the predictor variable coefficients from this model. A significant main effect of semantic dissimilarity prediction accuracy was found at the level of individual trials (t = 2.08, p = 0.04). These results indicate that an individual’s performance on the comprehension task overall and at the level of individual trials could be predicted from their neural responses to speech.

**Table 2.** Trial LME model. Model weights for the linear mixed effects model predicting comprehension scores at the level of individual trials.

|  | Coefficient | S.E. | t statistic | p |
| --- | --- | --- | --- | --- |
| **Sem. Dissim.** | **0.098** | **0.047** | **2.08** | **0.04** |
| Env. | 0.063 | 0.052 | 1.22 | 0.22 |
| Sem. Dissim. * Env. | - 0.013 | 0.044 | -0.27 | 0.77 |

## 4. Discussion

In this study we examined whether TRFs derived using semantic dissimilarity related specifically to the semantic processing of words following coherent, narrative context. For the unscrambled speech condition, we found reliable neural indices that were similar to those derived in previous experiments (Broderick et al., 2018). Like the previous findings, components of the TRF to unscrambled speech shared characteristics with the N400 ERP component. These neural indices of speech processing became weaker as soon as any scrambling of the speech signal was introduced. This pattern of neural responses was largely consistent with speech comprehension accuracy. As a group, the subjects tended to do better than 50%, which was not surprising given that they were hearing all of the words necessary to intuit some answers. However, at higher levels of scrambling their ability to meaningfully comprehend the narrative was significantly reduced and they failed to exceed the 95% confidence interval on the multiple choice comprehension questions. Meanwhile, lexical identification questionnaire scores were similar across scrambling conditions, allowing us to confidently rule out the possibility of large changes in subject engagement across scrambling conditions. Subjects were able to accurately identify words that appeared in a presented trial, indicating that they could recognise the words despite a drop in their overall understanding.

Our findings give further support to the idea that the semantic dissimilarity derived TRF reflects the semantic processing of words relative to their preceding context. When listening to speech, we incrementally build an internal representation of the current event, discourse, or context (Eberhard, Spivey-Knowlton, Sedivy, & Tanenhaus, 1995; W. Marslen-Wilson, 1973; W. D. Marslen-Wilson, 1975; Tanenhaus, Spivey-Knowlton, Eberhard, & Sedivy, 1995) that will influence how subsequent information is processed (Obleser, 2014). Such is the basis for language understanding. Our behavioural results are consistent with previous studies which observed a drop in behavioural performance with the introduction of scrambling (Bautista & Wilson, 2016; Humphries et al., 2006; Mollica et al., 2020). Together, these findings outline the importance of a coherent context for the comprehension of words and full speech excerpts. Crucially, our semantic dissimilarity measure was sensitive to speech comprehension, giving further support to the conclusion that the neural measure reflects contextual effects on language understanding (see Fig S1 (Broderick et al., 2018)) and refuting the recent suggestions that the semantic dissimilarity TRF may reflect the processing of content words more generally (Dijkstra et al., 2020). In addition, output measures from the semantic dissimilarity TRF were correlated with behavioural scores at the level of individual subjects and individual trials. These results represent a promising direction for future objective biomarkers of language comprehension that may be used in practical and clinical settings.

Unlike the semantic dissimilarity TRF, TRFs based on the speech envelope were unable to distinguish between scrambling conditions despite the behavioural difference in speech comprehension across conditions. Previous work has indicated that neural tracking of the speech envelope is foundational for speech comprehension (Peelle & Davis, 2012). However, our findings further highlight the fact that there is not a one-to-one mapping between the two measures (Brodbeck & Simon, 2020), and suggest that there is a limit to the stages of higher-level linguistic processing that envelope tracking measures can reliably index. The speech envelope is an important cue for intelligibility (Drullman et al., 1994a, 1994b; Shannon et al., 1995) and its clear from previous research that envelope tracking represents more than basic acoustic processing and is influenced by some speech-specific component (Peelle, Gross, & Davis, 2013; Prinsloo & Lalor, 2020). However, the speech envelope is a fundamentally low-level, impoverished representation of speech that measures amplitude fluctuations in acoustic energy. Although dominant frequencies in this signal correspond to the rate of important linguistic units on multiple time-scales (Ding et al., 2017), the represented linguistic features are conflated into a single univariate measure. Thus, it is unclear which features of speech are being tracked by the brain using the envelope-based TRF. In addition, dissociating the neural processes (for example acoustic processing, speech perception or speech comprehension) that underlie envelope tracking has been a major challenge (Ding & Simon, 2014). From our results, it seems clear however, that envelope tracking measures do not strongly index the context-based semantic processing that underlies natural speech comprehension. In contrast, TRFs based on semantic dissimilarity show a closer correspondence with behavioural measures of comprehension. This is unsurprising, given that the measure directly reflects the relationship between a word’s semantic features and those of its preceding words.

Previous work examining the relationship between envelope tracking and comprehension have found stronger tracking when speech is comprehended (Ahissar et al., 2001; Ding, Chatterjee, & Simon, 2014; Peelle et al., 2013; Vanthornhout, Decruy, Wouters, Simon, & Francart, 2018). However, these studies typically manipulate a listener’s understanding by changing the acoustics of the signal. Applying noise vocoding or lower SNR to a signal affects a listener’s ability to understand speech but also affects their ability to recognise words (Shannon et al., 1995). Therefore, as well as the confound of acoustically manipulating the signal, it has been challenging to dissociate processes of speech recognition and language comprehension using this type of experimental paradigm. Of course, neural indices of envelope tracking have provided great insight into how speech is processed from early acoustic stages (Ding et al., 2014; Ding & Simon, 2013; Drennan & Lalor, 2019) to stages of lexical identification (Ding, Melloni, Zhang, Tian, & Poeppel, 2016; Peelle et al., 2013). However, our results cast doubt on the utility of such measures as indices of higher-level processes such as comprehension. Researchers should take care when choosing this measure if their main goal is to investigate language understanding.

One additional observation for the envelope TRF was the significant (only when uncorrected) difference between weights at a time lag of ~200ms. This finding corresponds with results from Power and colleagues who estimated envelope TRF weights for individuals participating in a dichotic listening paradigm (Power, Foxe, Forde, Reilly, & Lalor, 2012). They found a late locus of selective attention at a time-lag of 200ms, where TRF weights for the attended speaker’s envelope were more positive than the unattended speaker’s envelope. The difference between weights for attended and unattended speakers in this study seems more pronounced than the difference between weights for our scrambling conditions. Furthermore, later studies using the same dataset could effectively distinguish between attended and unattended speakers using the TRF approach (O’Sullivan et al., 2015), whereas our envelope based TRF could not distinguish between scrambling conditions. Nevertheless, it’s possible that the differences between the envelope TRF weights across scrambling conditions in this window could reflect a filtering process located at the level of early semantic analysis (Treisman, 1964) that is better facilitated by higher contextual support. We remain circumspect on this though given that our effect did not survive correction for multiple comparisons. Future work should more directly investigate the relationship between output TRF measures and the speech representations they are based on.

Finally, while we have discussed the difficulties that come with acoustically modifying speech in order to affect its comprehensibility, the paradigm we have used here comes with its own challenges. Specifically, scrambling word order, as we have done, will not only disrupt the semantic flow of the narrative, but it will also profoundly disrupt the syntactic structure of the speech. As such, we cannot definitively rule out that syntactic processing may substantially contribute to our semantic dissimilarity measure, which would make it less of a pure measure of semantic speech comprehension than we have been arguing. However, we are somewhat sceptical of the idea of a large contribution from syntactic processing. The main reason for our scepticism is that syntactic processing has far more commonly been linked with the so-called P600 component (Kaan, Harris, Gibson, & Holcomb, 2000), a positive component that typically onsets around 500 ms and peaks at around 600 ms. The fact that the semantic dissimilarity TRF displays a much earlier negative peak makes us much more inclined to link it to semantic processing that has long been associated with the classic N400 component. That said, for years researchers have highlighted the challenges of dissociating semantic and syntactic processing in neurophysiological research, with some work linking syntactic processing to the N400 (Hagoort, 2003) and some suggesting semantic processing contributions to the P600 (Kuperberg, 2007). As such, we must remain at least somewhat circumspect in our interpretation of the semantic dissimilarity TRF. Future work presenting word lists with limited syntactic structure that still convey some form of narrative might help in further elucidating precisely what kinds of processing are indexed by this measure.

In conclusion, our results give further support that TRFs based on a semantic dissimilarity representation of speech reflect the processing of word meaning in context. The neural measure shows higher correspondence with speech comprehension than the much more commonly used measure of envelope tracking. We propose that the adoption of the relatively new semantic dissimilarity TRF approach may be beneficial for researchers wishing to investigate language processing using naturalistic stimuli.

## 5. Acknowledgements

This work was supported by a Career Development Award from Science Foundation Ireland (15/CDA/3316), an Irish Research Council Government of Ireland Postgraduate Scholarship (GOIPG/2015/3378), and the Del Monte Neuroscience Institute at the University of Rochester.

## References

Ahissar, E., Nagarajan, S., Ahissar, M., Protopapas, A., Mahncke, H., & Merzenich, M.M. (2001). Speech comprehension is correlated with temporal response patterns recorded from auditory cortex. Proceedings of the National Academy of Sciences, 98(23), 13367–13372.

Bautista, A., & Wilson, S. M. (2016). Neural responses to grammatically and lexically degraded speech. Language, Cognition and Neuroscience, 31(4), 567–574.

Benjamini, Y., & Hochberg, Y. (1995). Controlling the false discovery rate: a practical and powerful approach to multiple testing. Journal of the royal statistical society. Series B (Methodological), 289–300.

Brodbeck, C., & Simon, J. Z. (2020). Continuous speech processing. Current Opinion in Physiology.

Broderick, M. P., Anderson, A. J., Di Liberto, G. M., Crosse, M. J., & Lalor, E. C. (2018). Electrophysiological correlates of semantic dissimilarity reflect the comprehension of natural, narrative speech. Current Biology, 28(5), 803–809.

Broderick, M. P., Di Liberto, G. P., Anderson, A. J., Rofes, A., & Lalor, E. C. (2020). Dissociable electrophysiological measures of natural language processing reveal differences in comprehension strategy in healthy ageing. bioRxiv.

Combrisson, E., & Jerbi, K. (2015). Exceeding chance level by chance: The caveat of theoretical chance levels in brain signal classification and statistical assessment of decoding accuracy. Journal of Neuroscience Methods, 250, 126–136.

Davis, M. H., & Johnsrude, I. S. (2003). Hierarchical processing in spoken language comprehension. Journal of Neuroscience, 23(8), 3423–3431.

de Heer, W. A., Huth, A. G., Griffiths, T. L., Gallant, J. L., & Theunissen, F. E. (2017). The hierarchical cortical organization of human speech processing. Journal of Neuroscience, 3267–3216.

Di Liberto, G. M., O’Sullivan, J. A., & Lalor, E. C. (2015). Low-Frequency Cortical Entrainment to Speech Reflects Phoneme-Level Processing. Current Biology, 25(19), 2457–2465.

Dijkstra, K., Desain, P., & Farquhar, J. (2020). Exploiting Electrophysiological Measures of Semantic Processing for Auditory Attention Decoding. bioRxiv.

Ding, N., Chatterjee, M., & Simon, J. Z. (2014). Robust cortical entrainment to the speech envelope relies on the spectro-temporal fine structure. Neuroimage, 88, 41–46.

Ding, N., Melloni, L., Zhang, H., Tian, X., & Poeppel, D. (2016). Cortical tracking of hierarchical linguistic structures in connected speech. Nature Neuroscience, 19(1), 158–164. doi:10.1038/nn.4186

Ding, N., Patel, A. D., Chen, L., Butler, H., Luo, C., & Poeppel, D. (2017). Temporal modulations in speech and music. Neuroscience & Biobehavioral Reviews.

Ding, N., & Simon, J. Z. (2013). Adaptive Temporal Encoding Leads to a Background-Insensitive Cortical Representation of Speech. Journal of Neuroscience, 33(13), 5728–5735. doi:10.1523/jneurosci.5297-12.2013

Ding, N., & Simon, J. Z. (2014). Cortical entrainment to continuous speech: functional roles and interpretations. Frontiers in Human Neuroscience, 8.

Doelling, K. B., & Poeppel, D. (2015). Cortical entrainment to music and its modulation by expertise. Proceedings of the National Academy of Sciences, 112(45), E6233–E6242.

Drennan, D. P., & Lalor, E. C. (2019). Cortical tracking of complex sound envelopes: modeling the changes in response with intensity. eNeuro, ENEURO. 0082–0019.2019.

Drullman, R., Festen, J. M., & Plomp, R. (1994a). Effect of reducing slow temporal modulations on speech reception. The Journal of the Acoustical Society of America, 95(5), 2670–2680.

Drullman, R., Festen, J. M., & Plomp, R. (1994b). Effect of temporal envelope smearing on speech reception. The Journal of the Acoustical Society of America, 95(2), 1053–1064.

Eberhard, K. M., Spivey-Knowlton, M. J., Sedivy, J. C., & Tanenhaus, M. K. (1995). Eye movements as a window into real-time spoken language comprehension in natural contexts. Journal of Psycholinguistic Research, 24(6), 409–436.

Etard, O., & Reichenbach, T. (2019). Neural speech tracking in the theta and in the delta frequency band differentially encode clarity and comprehension of speech in noise. Journal of Neuroscience, 39(29), 5750–5759.

Frank, S. L., & Willems, R. M. (2017). Word predictability and semantic similarity show distinct patterns of brain activity during language comprehension. Language, Cognition and Neuroscience, 1–12.

Hagoort, P. (2003). Interplay between syntax and semantics during sentence comprehension: ERP effects of combining syntactic and semantic violations. Journal of Cognitive Neuroscience, 15(6), 883–899.

Hickok, G., & Poeppel, D. (2007). The cortical organization of speech processing. Nature Reviews Neuroscience, 8(5), 393–402.

Howard, M. F., & Poeppel, D. (2010). Discrimination of speech stimuli based on neuronal response phase patterns depends on acoustics but not comprehension. Journal of Neurophysiology, 104(5), 2500–2511.

Humphries, C., Binder, J. R., Medler, D. A., & Liebenthal, E. (2006). Syntactic and semantic modulation of neural activity during auditory sentence comprehension. Journal of Cognitive Neuroscience, 18(4), 665–679.

Hyvarinen, A. (1999). Fast and robust fixed-point algorithms for independent component analysis. IEEE Transactions on Neural Networks, 10(3), 626–634.

Kaan, E., Harris, A., Gibson, E., & Holcomb, P. (2000). The P600 as an index of syntactic integration difficulty. Language and Cognitive Processes, 15(2), 159–201.

Kiss, M., Cristescu, T., Fink, M., & Wittmann, M. (2008). Auditory language comprehension of temporally reversed speech signals in native and non-native speakers. Acta Neurobiologiae Experimentalis, 68(2), 204.

Kubanek, J., Brunner, P., Gunduz, A., Poeppel, D., & Schalk, G. (2013). The tracking of speech envelope in the human cortex. PloS One, 8(1), e53398.

Kuperberg, G. R. (2007). Neural mechanisms of language comprehension: Challenges to syntax. Brain Research, 1146, 23–49.

Kutas, M. (1993). In the company of other words: Electrophysiological evidence for single-word and sentence context effects. Language and Cognitive Processes, 8(4), 533–572.

Kutas, M., & Federmeier, K. D. (2011). Thirty years and counting: Finding meaning in the N400 component of the event related brain potential (ERP). Annual Review of Psychology, 62, 621.

Kutas, M., & Hillyard, S. A. (1980). Reading senseless sentences: Brain potentials reflect semantic incongruity. Science, 207(4427), 203–205.

Lalor, E. C., & Foxe, J. J. (2010). Neural responses to uninterrupted natural speech can be extracted with precise temporal resolution. European Journal of Neuroscience, 31(1), 189–193. doi:10.1111/j.1460-9568.2009.07055.x

Lalor, E. C., Power, A. J., Reilly, R. B., & Foxe, J. J. (2009). Resolving precise temporal processing properties of the auditory system using continuous stimuli. Journal of Neurophysiology, 102(1), 349–359. doi:10.1152/jn.90896.2008

Lerner, Y., Honey, C. J., Silbert, L. J., & Hasson, U. (2011). Topographic mapping of a hierarchy of temporal receptive windows using a narrated story. Journal of Neuroscience, 31(8), 2906–2915.

Luo, H., & Poeppel, D. (2007). Phase patterns of neuronal responses reliably discriminate speech in human auditory cortex. Neuron, 54(6), 1001–1010.

Marslen-Wilson, W. (1973). Linguistic structure and speech shadowing at very short latencies. Nature.

Marslen-Wilson, W. D. (1975). Sentence perception as an interactive parallel process. Science, 189(4198), 226–228.

Mollica, F., Siegelman, M., Diachek, E., Piantadosi, S. T., Mineroff, Z., Futrell, R., & Fedorenko, E. (2018). High local mutual information drives the response in the human language network. bioRxiv, 436204.

Mollica, F., Siegelman, M., Diachek, E., Piantadosi, S. T., Mineroff, Z., Futrell, R., … Fedorenko, E. (2020). Composition is the core driver of the language-selective network. Neurobiology of Language, 1(1), 104–134.

Naselaris, T., Kay, K. N., Nishimoto, S., & Gallant, J. L. (2011). Encoding and decoding in fMRI. Neuroimage, 56(2), 400–410.

O’Sullivan, J. A., Power, A. J., Mesgarani, N., Rajaram, S., Foxe, J. J., Shinn-Cunningham, B. G., … Lalor, E. C. (2015). Attentional selection in a cocktail party environment can be decoded from single-trial EEG. Cerebral cortex, 25(7), 1697–1706.

Obleser, J. (2014). Putting the listening brain in context. Language and linguistics compass, 8(12), 646–658.

Obleser, J., Herrmann, B., & Henry, M. J. (2012). Neural oscillations in speech: don’t be enslaved by the envelope. Frontiers in Human Neuroscience, 6, 250.

Oord, A. v. d., Dieleman, S., Zen, H., Simonyan, K., Vinyals, O., Graves, A., … Kavukcuoglu, K. (2016). Wavenet: A generative model for raw audio. arXiv preprint arXiv:1609.03499.

Peelle, J. E., & Davis, M. H. (2012). Neural oscillations carry speech rhythm through to comprehension. Frontiers in Psychology, 3.

Peelle, J. E., Gross, J., & Davis, M. H. (2013). Phase-locked responses to speech in human auditory cortex are enhanced during comprehension. Cerebral cortex, 23(6), 1378–1387.

Peña, M., & Melloni, L. (2012). Brain oscillations during spoken sentence processing. Journal of Cognitive Neuroscience, 24(5), 1149–1164.

Pennington, J., Socher, R., & Manning, C. D. (2014). Glove: Global vectors for word representation. Paper presented at the Proceedings of the 2014 conference on empirical methods in natural language processing (EMNLP).

Poeppel, D., Emmorey, K., Hickok, G., & Pylkkänen, L. (2012). Towards a new neurobiology of language. Journal of Neuroscience, 32(41), 14125–14131.

Power, A. J., Foxe, J. J., Forde, E. J., Reilly, R. B., & Lalor, E. C. (2012). At what time is the cocktail party? A late locus of selective attention to natural speech. European Journal of Neuroscience, 35(9), 1497–1503. doi:10.1111/j.1460-9568.2012.08060.x

Price, C. J. (2010). The anatomy of language: a review of 100 fMRI studies published in 2009. Annals of the New York Academy of Sciences, 1191(1), 62–88.

Prinsloo, K. D., & Lalor, E. C. (2020). General auditory and speech-specific contributions to cortical envelope tracking revealed using auditory chimeras. bioRxiv.

Saberi, K., & Perrott, D. R. (1999). Cognitive restoration of reversed speech. Nature, 395(6730), 760–760.

Shannon, R. V., Zeng, F.-G., Kamath, V., Wygonski, J., & Ekelid, M. (1995). Speech recognition with primarily temporal cues. Science, 270(5234), 303–304.

Tanenhaus, M. K., Spivey-Knowlton, M. J., Eberhard, K. M., & Sedivy, J. C. (1995). Integration of visual and linguistic information in spoken language comprehension. Science, 1632–1634.

Treisman, A. M. (1964). Verbal cues, language, and meaning in selective attention. The American journal of psychology, 206–219.

Vanthornhout, J., Decruy, L., Wouters, J., Simon, J. Z., & Francart, T. (2018). Speech intelligibility predicted from neural entrainment of the speech envelope. Journal of the Association for Research in Otolaryngology, 19(2), 181–191.

Verschueren, E., Somers, B., & Francart, T. (2019). Neural envelope tracking as a measure of speech understanding in cochlear implant users. Hearing Research, 373, 23–31.

White, E.B. (1951) Charlotte’s Web. Harper and Brothers.

